# Human relevant exposure to di-n-butyl phthalate tampers with the ovarian insulin-like growth factor 1 system and disrupts folliculogenesis in young adult mice

**DOI:** 10.1101/2023.03.15.532792

**Authors:** Estela J. Jauregui, Maile McSwain, Xiaosong Liu, Kara Miller, Kimberlie Burns, Zelieann R. Craig

## Abstract

Phthalates are compounds used in consumer and medical products worldwide. Phthalate exposure in women has been demonstrated by detection of phthalate metabolites in their urine and ovarian follicular fluid. High urinary phthalate burden has been associated with reduced ovarian reserve and oocyte retrieval in women undergoing assisted reproduction. Unfortunately, no mechanistic explanation for these associations is available. In short term in vivo and in vitro animal studies modeling human relevant exposures to di-n-butyl phthalate (DBP), we have identified ovarian folliculogenesis as a target for phthalate exposures. In the present study, we investigated whether DBP exposure negatively influences insulin-like growth factor 1 (IGF) signaling in the ovary and disrupts ovarian folliculogenesis. CD-1 female mice were exposed to corn oil (vehicle) or DBP (10 or 100 μg/kg/day) for 20-32 days. Ovaries were collected as animals reached the proestrus stage to achieve estrous cycle synchronization. Levels of mRNAs encoding IGF1 and 2 (*Igf1* and *Igf2*), IGF1 receptor (*Igf1r*), and IGF binding proteins 1-6 (*Ifgbp1-6*) were measured in whole ovary homogenates. Ovarian follicle counts and immunostaining for phosphorylated IGF1R protein (pIGF1R) were used to evaluate folliculogenesis and IGF1R activation, respectively. DBP exposure, at a realistic dose that some women may experience (100 μg/kg/day for 20-32 days), reduced ovarian *Igf1* and *Igf1r* mRNA expression and reduced small ovarian follicle numbers and primary follicle pIGF1R positivity in DBP-treated mice. These findings reveal that DBP tampers with the ovarian IGF1 system and provide molecular insight into how phthalates could influence the ovarian reserve in females.

## INTRODUCTION

Phthalates are a family of highly versatile chemicals used in the manufacture of a variety of products ranging from personal care and consumer products to medical devices and medication coatings. Their detection in a variety of human biofluids and their associations with several adverse reproductive health outcomes support their designation as endocrine-disrupting chemicals. Specifically, phthalates have been detected in human urine, amniotic fluid, and ovarian follicular fluid (Calafat *et al.*, 2006; M. J. Silva *et al.*, 2004; Manori J Silva *et al.*, 2004; Du *et al.*, 2016) and associated with increased risk for early menopause, low ovarian reserve, and low egg retrieval in women (Grindler *et al.*, 2015; Messerlian *et al.*, 2016; Hauser *et al.*, 2016). Unfortunately, the mechanisms underlying these associations are not understood and require further evaluation using experimental models.

Among phthalates, di-n-butyl phthalate (DBP) is of particular interest based on its use in personal care products and medications, and higher levels of its metabolite, mono-n-butyl phthalate (MBP), in females (Centers for Disease Control and Prevention, 2013). Despite knowledge of DBP exposure estimates for the general population (7-10 μg/kg/day; (Kavlock *et al.*, 2002)), occupational settings (0.1-76 μg/kg/day; (Hines *et al.*, 2011)), and coated medication ingestion (1-233 μg/kg/day; (Hernández-Díaz *et al.*, 2013)), very few animal studies have evaluated the reproductive toxicity of DBP at low, human relevant doses. To address this gap, we have used cycling CD-1 female mice treated with human relevant doses to demonstrate that oral DBP exposure leads to detection of MBP in the ovary (Jauregui *et al.*, 2021), reduces diestrus serum 17β-estradiol (E2) concentration (Sen *et al.*, 2015), reduces diestrus antral follicle counts (Sen *et al.*, 2015), and impairs ovarian gene expression (Sen *et al.*, 2015; Liu and Craig, 2019; Jauregui *et al.*, 2021).

Folliculogenesis depends on the interplay of several hormones, including follicle-stimulating hormone (FSH), E_2_, and insulin-like growth factor (IGF). The IGF system is comprised of two insulin-like growth factors, IGF1 and IGF2, two insulin-like growth factor receptors, IGF1R and IGF2R, and six IGF binding proteins (IGFBPs) which regulate the availability of IGF1 (Annunziata *et al.*, 2011). The ovary has a local IGF system previously shown to be an essential paracrine pathway for the growth and selection of ovarian follicles in several species (Mazerbourg *et al.*, 2003). Although species differences in the relative importance of each IGF ligand exist (IGF1 for rodents, IGF2 for primates), the conserved IGF1R receptor localization and function in various species (Bondy *et al.*, 1993) supports the use of rodent models to evaluate toxicity to this system experimentally while retaining relevance to humans. Genetic or pharmacological inhibition of IGF1R signaling in female mice results in a lack of antral follicle progression to pre-ovulatory stage, reduced serum E_2_ concentration, impaired FSH action in the ovary, and overall infertility (Baumgarten *et al.*, 2017; Zhou *et al.*, 1997). Interestingly, all four of these phenotypes are similar to outcomes observed in association with phthalate exposures in various studies (Sen *et al.*, 2015; Messerlian *et al.*, 2016; Wang *et al.*, 2016) and reviewed recently (Land *et al.*, 2022).

While the IGF system has been recognized as vulnerable to endocrine disruption (Talia *et al.*, 2021), knowledge regarding the effects of endocrine-disrupting chemicals on gonadal IGF systems is currently limited to prenatal exposures to 2,3,7,8-Tetrachlorodibenzo-p-dioxin in female rats (Zhang *et al.*, 2019), di(2-ehtylhexyl) phthalate in male rats (DEHP; (Lin *et al.*, 2008), and bisphenol A in the KGN granulosa cell line (Kwintkiewicz *et al.*, 2010). Human epidemiological studies have reported an inverse relationship between some phthalates and the different components of the systemic IGF system in humans (Wu *et al.*, 2017; Watkins *et al.*, 2016; Zhao *et al.*, 2016; Montrose *et al.*, 2018; LaRocca *et al.*, 2014) however, no studies have investigated the influence of phthalates on the ovarian IGF system following direct oral exposure. Therefore, in the present study, we use CD-1 female mice orally exposed to human relevant doses of DBP to investigate whether DBP exposure influences the ovarian IGF system and disrupts folliculogenesis.

## MATERIALS AND METHODS

### Animals

All animal experiments were performed as described following the guidelines stated in the Guide for the Care and Use of Laboratory Animals (National Research Council (U.S.) *et al.*, 2011). All animal work was approved by the Institutional Animal Care and Use Committee at the University of Illinois, where the experiments were conducted.

Young adult (28 days old) female CD-1 mice were obtained from Charles River Laboratories (Charles River, California) and housed in single-use BPA-free cages at the University of Illinois College of Veterinary Medicine Central Animal Facility (Z.R. Craig’s previous institution). Upon arrival at the animal facility, mice were allowed to acclimate for at least 24 h before handling. Water and food were provided *ad libitum*, room cycles set to 12 L:12 D cycles, and temperature maintained at 22 ± 1°C. Mice were subjected to daily weighing, vaginal smear collections, and oral dosing with vehicle or DBP treatments. At the end of each study, mice were euthanized under pre-sedation by carbon dioxide (CO_2_) inhalation followed by cervical dislocation.

### Chemicals

Tocopherol-stripped corn oil was obtained from MP Biomedicals (Solon, OH). Dibutyl phthalate (DBP, CAS No. 84-74-2; 99.6% purity) was obtained from Sigma-Aldrich (St Louis, Missouri).

### Dosing and tissue collection

Female CD-1 mice (n=16/treatment, 35 days old) were randomly assigned to receive oral tocopherol-stripped corn oil (vehicle) or one of three DBP dose levels that were inspired by human daily estimates (10 μg/kg/day and 100 μg/kg/day) or classical toxicity testing high doses (1000 mg/kg/day) as previously described (Sen *et al.*, 2015). Animals were subjected to vaginal smears, weighed, and dosed daily for at least 20 consecutive days and then euthanized as they reached the proestrus stage. The selected dosing and collection scheme resulted in proestrus-matched mice that were dosed for 20-32 days. At the end of the experiment, ovaries were dissected from each animal and rid of fat and oviductal tissue. One ovary from each cleaned pair was snap-frozen for RNA extraction, while the other was fixed for subsequent histological processing.

### Ovarian follicle counts

Ovaries were fixed in 10% formalin overnight at 4°C. After fixing, ovaries were washed in 70% ethanol, processed, and embedded in paraffin. Embedded ovaries were sectioned (5 μm thickness) and processed for hematoxylin (Richard-Allan, 7211) and eosin (Richard-Allan, 7111) staining. Stained ovary slides were subjected to follicle classification and enumeration procedures as previously described (Sen *et al.*, 2015; Liu and Craig, 2019). Briefly, follicles containing visible oocytes were counted blinded on every 20^th^ section of the ovary and classified as primordial (oocyte surrounded by a layer of squamous granulosa cells), primary (oocyte surrounded by a layer containing at least half of cuboidal granulosa cells), secondary (oocyte surrounded by at least two layers of cuboidal granulosa cells and theca cells layer), or antral (oocyte surrounded by layers of cuboidal granulosa cells and theca cells, and an antrum formed). Additionally, follicles were classified as atretic if they had at least one of the following: (1) follicular cell pyknosis, (2) granulosa/theca cell layer disorganization, (3) oocyte fragmentation, and (4) theca cell hypertrophy (Liu and Craig, 2019). Follicles that were not atretic but showed abnormal features (i.e., missing cells, abnormal shape) were classified as abnormal.

### RNA extraction and cDNA synthesis

Snap frozen ovaries were subjected to RNA extraction with DNAse treatment using Qiagen RNeasy Micro Kits (Qiagen, Valencia, California). RNA concentrations were determined at 260nm using a Take3 microvolume plate on a Synergy H1m microplate reader (Biotek, Winooski, Vermont). RNA samples (1 μg) were reverse transcribed using iScript cDNA synthesis kits (Bio-Rad, Hercules, California) to generate cDNA for qPCR.

### Quantitative Polymerase Chain Reaction (qPCR)

Quantitative PCR was performed for each sample using 1 μl of cDNA (0.5 μg), 1 μl of each primer (5 μM), 5 μl of Ssofast EvaGreen Supermix (Bio-Rad, Hercules, California), and 2 μl of nuclease-free water. Each reaction was performed in triplicate, and no template, no primer, and no reverse transcriptase controls were included. Primers used for *Igf1* (forward: 5’-ATCCCTTCCAACCAGTGGCTGACC-3’ and reverse: 5’-GGAGCCTCCTGCCAAGTGTTTAGC-3’), *Igf2* (forward: 5’-CATCGTCCCCTGATCGTGTTAC-3’ and reverse: 5’-GGAACTGTCCCTGCTCAAGA-3’), *Igf1r* (forward: 5-AGCAAGTTCTTCGTTTCGTCA-3’ and reverse: 5’-CTCCATCTCATCCTTGATGCT-3’), *Igf1bp1* (forward: 5’-CCGACCTCAAGAAATGGAA-3’ and reverse: 5’-CATCTCCTGCTTTCTGTTGG-3’), *Igf1bp2* (forward: 5’-ATCTCTACTCCCTGCACATCC-3’ and reverse: 5’-TCCGTTCAGAGACATCTTGC-3’), *Igf1bp3* (forward: 5’-CACATCCCAAACTGTGACAA-3’ and reverse: 5’-CCATACTTGTCCACACACCA-3’), *Igf1bp4* (forward: 5’-ATCCCCATTCCAAACTGTGA-3’ and reverse: 5’-GATCCACACACCAGCACTTG-3’), *Igf1bp5* (forward: 5’-ACTGTGACCGCAAAGGATTC-3’ and reverse: 5’-TTGTCCACACACCAGCAGAT-3’), *Igf1bp6* (forward: 5’-AGAGGCTTCTACCGAAAGCA-3’ and reverse: 5’-TCCTTGACCATCTGGAGACA-3’), *Tbp* (forward: 5’-GTGCCAGATACATTCCGCCT-3’ and reverse: 5’-AGCTGCGTTTTTGTGCAGAG-3’), and *Actb* (forward: 5’-ATGCCGGAGCCGTTGTC-3’ and reverse: 5’-GCGAGCACAGCTTCTTTG-3’) were based on published sequences (Okano and Kelley, 2013; Sen *et al.*, 2015) and verified by PrimerBLAST software. The reference genes were confirmed to not be differentially expressed between treatments prior to analysis of the genes of interest. Data were analyzed using the ΔΔCt model for relative quantification and normalized to the average of the housekeeping genes, *Actb* and *Tbp*.

### Immunohistochemistry

Fixed ovaries were processed and embedded in paraffin. Ovarian sections (5 μm) were placed on charged slides, deparaffinized and rehydrated to distilled water. Antigen retrieval was performed using citrate buffer (Vector Labs H-3300; 30 min in steamer) followed by TBS (0.05M Tris buffered saline, Alfa aesar AAJ6076K2) buffer wash, peroxidase quenching with 0.5% H_2_O_2_ (20 min at room temperature), TBST (TBS + 0.5% Tween 20) buffer wash, universal blocker (Vector Labs SP-5035-100; 1 h), avidin block (Vector Labs SP-2001; 15 min), TBST buffer wash, and biotin block (Vector Labs SP-2001; 15 minutes). Slides were then incubated with primary anti-phosphorylated IGF1R antibody (Cat#3021, Cell Signaling Technology Inc.; Danvers, MA) at 2.11 μg/ml or protein concentration matched Isotype (overnight at 4°C). After washing in TBST, slides were incubated with biotinylated secondary (Vector Labs BP-9100-500; 1 h) followed by TBST wash, avidin-biotin complex-horseradish peroxidase (Vector labs PK-6100; 40 min), TBS wash, and DAB chromogen with nickel (Vector Labs SK-4100; 6 min). After a distilled water wash, counterstaining was performed using Hematoxylin QS (Vector Labs H-3403; 20 sec), followed by a distilled water wash, Bluing reagent (Richard-Allan #7301; 30 sec), another distilled water wash, dehydration to xylene and finally cover slipping with DPX (Electron Microscopy Sciences, #13512). All steps were performed at room temperature unless otherwise indicated. All washes were performed 3 times for 3 minutes each. Sections of tissue from mouse embryos (embryonic day 9.5) were used as a positive control, and rabbit IgG isotype (Cat#10500C, Invitrogen; Waltham, MA) was used as a negative control. At least two sections per ovary were selected for quantification. Sections were at least 100 μm from the ovary’s heel and 100 μm apart from each other. Counted follicles were then further classified as positive or negative based on observed immunostaining in the granulosa cells or surrounding the oocyte. A total of three ovaries per treatment group were analyzed.

### Statistical analyses

GraphPad Prism software (version 9.1) was used for all statistical analyses. All data were subjected to normality and homogeneity of variance test (Shapiro-Wilk) *a priori* to determine appropriate statistical tests. Parametric data were analyzed using one-way ANOVA followed by Dunnet’s *post hoc test.* When normally distributed data failed to meet the homogeneity of variances requirement, Welch’s ANOVA followed by Dunnett’s T3 multiple comparisons test was used instead. If a trend was observed between control and a phthalate group, values were further compared using Sidak’s multiple comparisons test. ROUT (Q=5%) was used to test whether any extreme values were statistically significant outliers. Non-parametric data were compared using the Kruskal-Wallis non-parametric test, followed by Dunn’s multiple comparisons test. Statistical significance was assigned at p≤0.05 for all comparisons.

## RESULTS

### Effect of DBP exposure on insulin-like growth factors and type 1 receptor mRNA levels

Levels of mRNAs encoding *Igf1, Igf2*, and *Igf1r* were detected in whole ovary homogenates from mice treated with oil or DBP and collected in proestrus. In comparison to vehicle treatment, exposure to DBP at 100 μg/kg/day significantly decreased the levels of *Igf1* (**Figure 1A**, p=0.028) and *Igf1r* (**Figure 1C**, p=0.048) without affecting *Igf2* (**Figure 1B,** p=0.997) which, in agreement with previous studies (Wandji *et al.*, 1998), showed lower abundance (average Ct>30) versus *Igf1.* The mRNA levels for all three genes did not differ from controls in the other DBP treatment groups tested (DBP 10 μg/kg/day and 1000 mg/kg/day).

**Figure 1.**
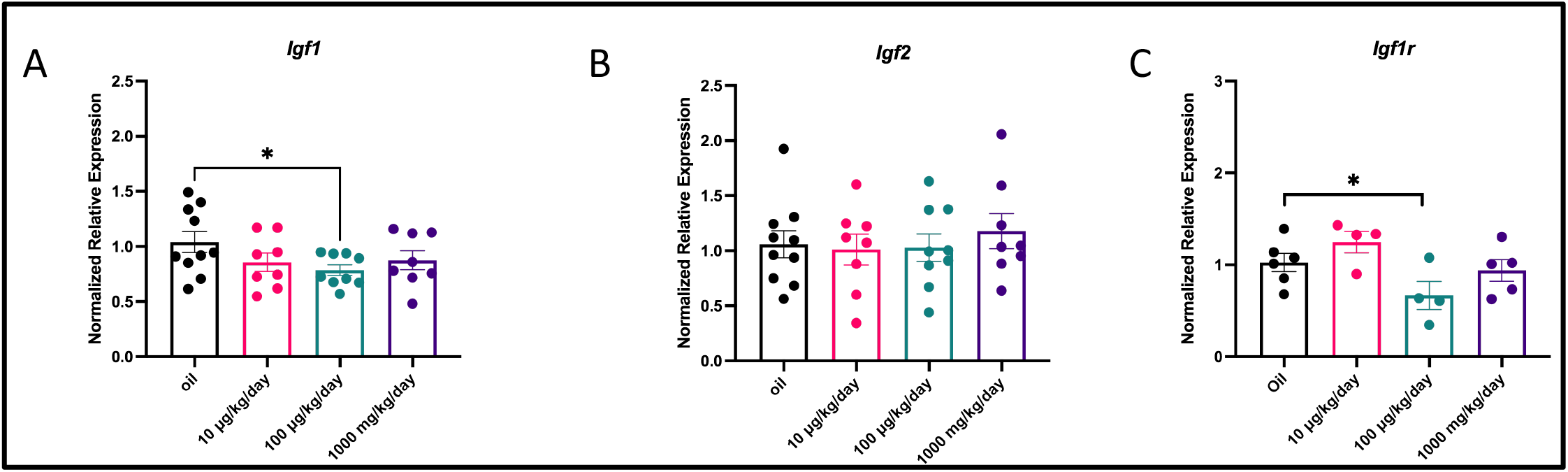
Effect of oral DBP exposure on ovarian insulin-like growth factors and type 1 receptor mRNA. Young adult cycling female CD-1 mice were treated daily with vehicle (oil) or DBP and their ovaries processed for qPCR as described in Materials and Methods. The expression of *Igf1* (**A**), *Igf2* (**B**), and *Igf1r* (**C**) was normalized to housekeeping genes *Actb* and *Tbp.* Data are presented as mean normalized relative expression ± SEM. Asterisks (*) indicate statistical significance at the p ≤ 0.05 level (n=8-10 mice/treatment).

### Effect of DBP exposure on insulin-like growth factor binding protein mRNA levels

Levels of mRNAs encoding *Igfbp1* through *6* were detected in whole ovary homogenates from oil- and DBP-treated mice. In comparison to vehicle treatment, exposure to DBP did not cause significant deviations in the expression of transcripts encoding IGF binding proteins (**Table 1**).

**Table 1.** Effect of oral DBP exposure on insulin-like growth factor binding proteins 1-6 mRNAs.

| Gene Name | Symbol | Treatment | Normalized Expression | Sample Number | Adj. P-value (vs. oil) |
| --- | --- | --- | --- | --- | --- |
| insulin-like growth factor binding protein 1 | *Igfbp1* | oil | 1.07 ± 0.12 | 10 | N/A |
|  |  | DBP 10µg/kg/day | 0.79 ± 0.10 | 8 | 0.4008 |
|  |  | DBP 100µg/kg/day | 0.86 ± 0.15 | 9 | 0.5997 |
|  |  | DBP 1000mg/kg/day | 1.20 ± 0.21 | 8 | 0.8822 |
| insulin-like growth factor binding protein 2 | *Igfbp2* | oil | 1.05 ± 0.11 | 10 | N/A |
|  |  | DBP 10µg/kg/day | 1.05 ± 0.12 | 8 | >0.9999 |
|  |  | DBP 100µg/kg/day | 1.06 ± 0.12 | 9 | 0.9999 |
|  |  | DBP 1000mg/kg/day | 0.97 ± 0.06 | 8 | 0.9329 |
| insulin-like growth factor binding protein 3 | *Igfbp3* | oil | 1.04 ± 0.09 | 10 | N/A |
|  |  | DBP 10µg/kg/day | 0.93 ± 0.15 | 8 | 0.8520 |
|  |  | DBP 100µg/kg/day | 0.94 ± 0.11 | 9 | 0.8697 |
|  |  | DBP 1000mg/kg/day | 0.84 ± 0.09 | 8 | 0.4817 |
| insulin-like growth factor binding protein 4 | *Igfbp4* | oil | 1.10 ± 0.16 | 10 | N/A |
|  |  | DBP 10µg/kg/day | 1.05 ± 0.13 | 8 | >0.9999 |
|  |  | DBP 100µg/kg/day | 1.24 ± 0.17 | 9 | >0.9999 |
|  |  | DBP 1000mg/kg/day | 1.11 ± 0.17 | 8 | >0.9999 |
| insulin-like growth factor binding protein 5 | *Igfbp5* | oil | 1.05 ± 0.10 | 10 | N/A |
|  |  | DBP 10µg/kg/day | 0.96 ± 0.12 | 8 | 0.8853 |
|  |  | DBP 100µg/kg/day | 1.12 ± 0.12 | 9 | 0.9377 |
|  |  | DBP 1000mg/kg/day | 0.98 ± 0.12 | 8 | 0.9525 |
| insulin-like growth factor binding protein 6 | *Igfbp6* | oil | 1.05 ± 0.11 | 10 | N/A |
|  |  | DBP 10µg/kg/day | 0.81 ± 0.08 | 8 | 0.1202 |
|  |  | DBP 100µg/kg/day | 0.90 ± 0.06 | 9 | 0.4748 |
|  |  | DBP 1000mg/kg/day | 0.81 ± 0.05 | 8 | 0.128 |

### Effect of DBP exposure on ovarian follicle numbers

Ovarian sections were subjected to follicle classification and enumeration to determine the impact of environmentally relevant DBP exposure on ovarian folliculogenesis. The total number of ovarian follicles counted per ovary was reduced in mice treated with DBP at 100 μg/kg/day in comparison to control (VEH: 278.5 ± 24.267; DBP100μg: 203.2 ± 22.296; p=0.02; **Figure 2A**). There were no significant differences in the number of total follicles per ovary in the 10 μg/kg/day or 1000 mg/kg/day DBP treatment groups compared to control (**Figure 2A**). Differential counts by follicle developmental stage revealed significant changes in primordial and primary follicle counts. Specifically, significantly low primordial follicle counts were observed in mice treated with DBP at 100 μg/kg/day (61.5 ± 7.6 follicles/ovary; p=0.006) and 1000 mg/kg/day (63 ± 4.0 follicles/ovary; p=0.02) compared to oil-treated controls (96.2 ± 9.6; **Figure 2B**). Furthermore, fewer primary follicles were observed in ovaries from mice treated with DBP at 100 μg/kg/day when compared to oil-treated mouse ovaries (VEH: 64.3 ± 8.8 follicles/ovary; DBP100μg: 44.3 ± 5.5 follicles/ovary; p=0.04; **Figure 2C**). There were no significant differences between the control group and the 10 μg/kg/day group for primordial and primary follicles or between control and the 1000 mg/kg/day group for primary follicles (**Figures 2B and 2C**). Similarly, the number of secondary follicles (**Figure 2D**), antral follicles (**Figure 2E**), atretic and abnormal follicles, or the percentage of total healthy and unhealthy follicles did not differ between treatments (data not shown).

**Figure 2.**
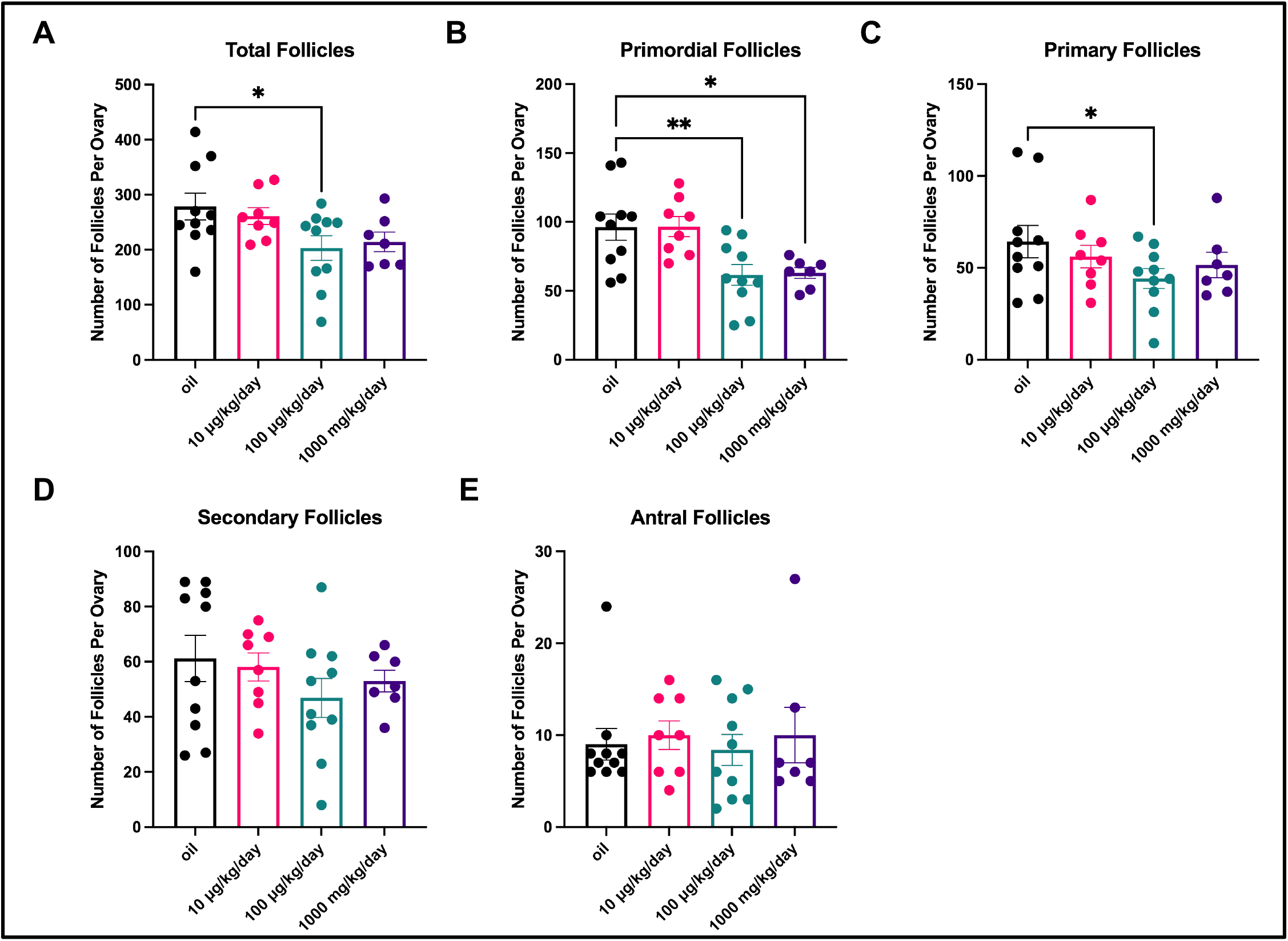
Effect of oral DBP exposure on ovarian follicle counts.

Young adult cycling female CD-1 mice were treated daily with vehicle (oil) or DBP and their ovaries processed for H&E staining and follicle counting as described in Materials and Methods. Total follicles (**A**), primordial follicles (**B**), primary follicles (**C**), secondary follicles (**D**), and antral follicles (**E**) from one ovary per mouse were classified and quantified. Data are presented as mean ± SEM. Asterisks indicate statistical significance at the p ≤ 0.05 (*) and p ≤ 0.01 (**) levels (n=7-10 mice/treatment).

### Localization of phosphorylated IGF1R protein immunostaining in the ovary

Immunohistochemical staining for an IgG matched to the anti-IGF1R antibody showed no staining (**Figure 3A;** negative control). Phosphorylated IGF1R (pIGF1R) immunostaining was observed in the granulosa cell compartment and surrounding the oocytes of primary, secondary, and antral follicles regardless of treatment (**Figures 3B-F**). Interestingly, pIGF1R immunostaining was also observed in atretic follicles (**Figure 3F**) and in co-localization with zona pellucida remnants in afollicular oocytes from presumed atretic follicles (not shown).

**Figure 3.**
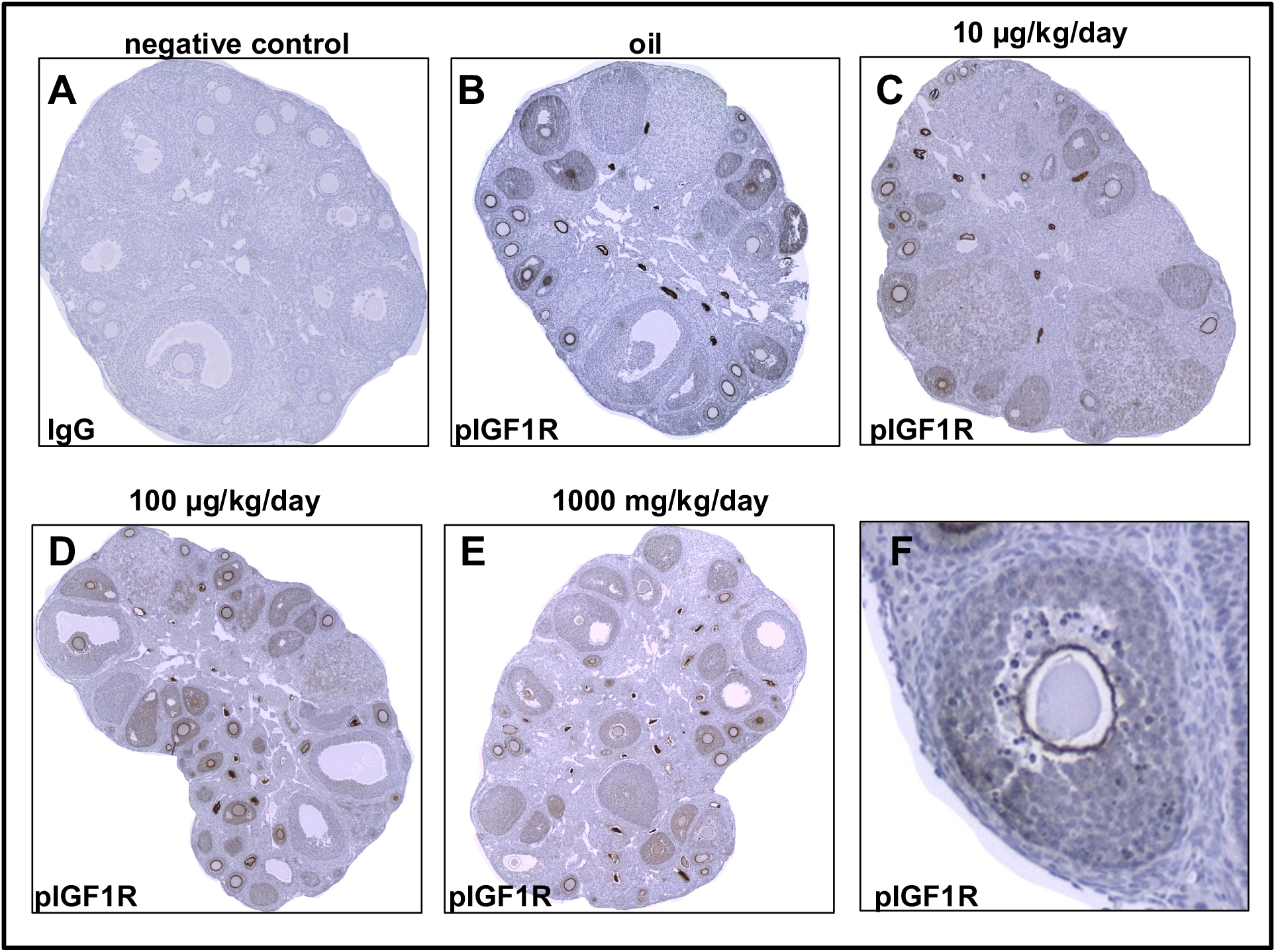
Localization of phosphorylated IGF1R (pIGF1R) protein in the ovaries of mice exposed to oil or DBP at various doses. Young adult cycling female CD-1 mice were treated daily with vehicle (oil) or DBP and their ovaries processed for immunohistochemical staining with anti-pIGF1R and its matched IgG (negative control) as described in Materials and Methods. Representative images from ovarian sections show: no staining for the antibody protein-matched isotype negative control (**A**), expression of IGF1R (depicted by brown staining) in vehicle control (**B**) or DBP 10 μg/kg/day (**C**), 100 μg/kg/day (**D**), and 1000 mg/kg/day (**E**) ovaries. Positive staining was observed in atretic follicles (**F**).

Evaluation of follicular pIGF1R positivity across the different follicular stages in oil-treated mice revealed a lack of pIGF1R in primordial follicles, with its detection starting at the primary follicle stage. Most pIGF1R-positive follicles observed were in the secondary and small antral stage (**Figure 4A-B**). Finally, quantification of pIGF1R-positive large antral follicles was attempted but found to be highly variable due to the low numbers observed (data not shown).

**Figure 4.**
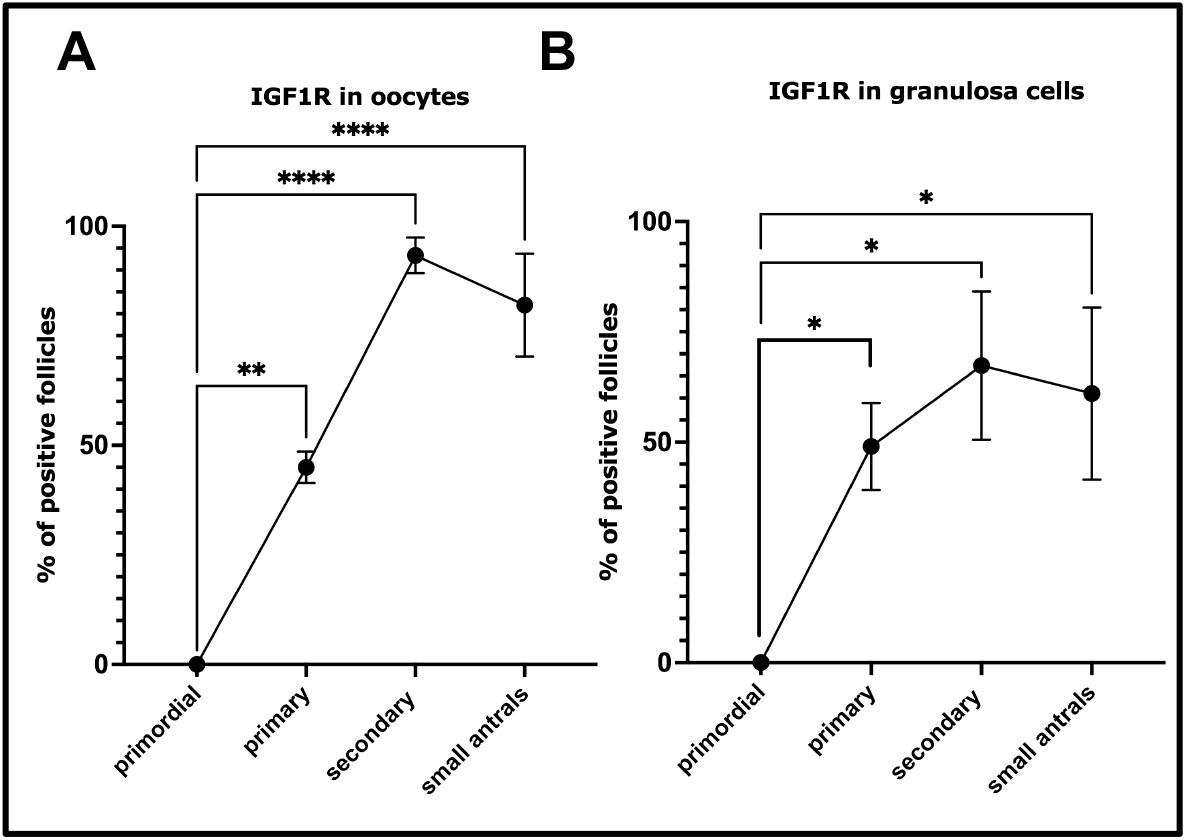
Percentage of phosphorylated IGF1R (pIGF1R)-positive follicles by developmental stage in control ovaries. Ovaries processed for immunohistochemical staining with anti-pIGF1R as described in Materials and Methods. At least two sections were evaluated to determine the total number and developmental stage of follicles per mouse ovary. Counted follicles were then further classified as positive or negative based on immunostaining in the granulosa cells or surrounding the oocyte. Data are presented as mean ± SEM. Asterisks indicate statistical significance at the p ≤ 0.05 (*), p ≤ 0.01 (**), and p ≤ 0.0001 (****) levels (n=3 mice/treatment).

### Effect of DBP exposure on the proportion of pIGF1R-positive ovarian follicles

Immunostaining for pIGF1R was evaluated based on the observation that DBP exposure resulted in significantly reduced *Igf1r* mRNA and primordial and primary follicle numbers in treated mice. The total number of follicles present and the number of pIGF1R-positive follicles (by oocyte and granulosa cells) were determined for stained ovarian sections from oil- and DBP-treated mice. The total percentage of pIGF1R-positive oocyte follicles was significantly decreased in mouse ovaries treated with DBP at 100 μg/kg/day when compared to vehicle (VEH: 51.33 ± 4.67%; DBP100μg: 32.67 ± 3.93%; p=0.03; **Figure 5A)**. Only a trend for reduced positive follicles was observed in the DBP 100 μg/kg/day group when considering pIGF1R immunostaining in granulosa cells only (VEH: 41.97 ± 0.12%; DBP100μg: 25.67 ± 3.38%; p=0.053; **Figure 5B**). Differential counts by follicle type revealed that differences in the proportion of pIGF1R positive follicles were driven by reductions in primary follicle positivity (**Figure 5C-D**). Interestingly, no differences in pIGF1R positive follicles were observed between vehicle and mice treated with DBP at the other doses (10 μg/kg/day and 1000 mg/kg/day).

**Figure 5.**
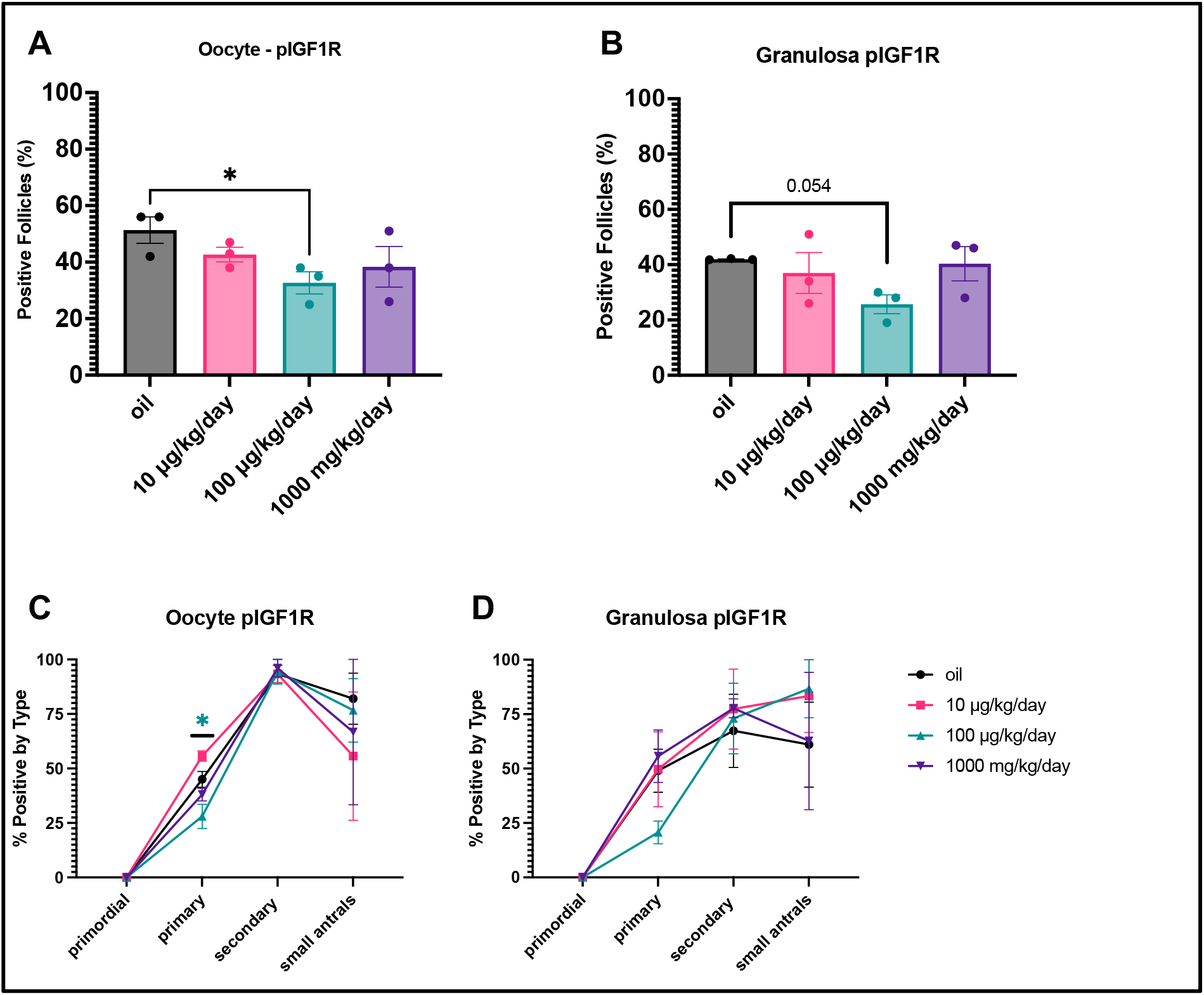
Effect of oral DBP exposure on the percentage of phosphorylated IGF1R (pIGF1R)-positive follicles. Ovaries were processed for immunohistochemical staining with anti-pIGF1R as described in Materials and Methods. At least two sections were evaluated to determine the total number and developmental stage of follicles per mouse ovary. Counted follicles were then further classified as positive or negative based on immunostaining in the granulosa cells or surrounding the oocyte. Data are presented as mean ± SEM. Asterisks indicate statistical significance at the p ≤ 0.05 (*) level (n=3 mice/treatment).

## DISCUSSION

In this study, young adult CD-1 mice were exposed to two environmentally relevant (10 and 100 μg/kg/day) and one classical high dose level (1000 μg/kg/day) of DBP to investigate whether phthalate exposure negatively influences ovarian IGF1 signaling components and ovarian folliculogenesis. Our findings show that DBP, at a dosage that occupationally and medication-exposed women could experience over their reproductive cycle (100 μg/kg/day for 20-32 days), results in dysregulated IGF1 signaling and reduced small ovarian follicle counts in mice. We did not detect any differences in the expression of mRNAs encoding IGF binding proteins, or in the number of secondary, antral, and unhealthy follicles between treatments. Taken together, our findings using a mouse model exposed to human relevant levels of DBP reveal that this phthalate may reduce the availability of ovarian IGF signaling components and, therefore, influence the ovarian reserve.

The mouse ovary has a complete intraovarian IGF1 system (Adashi *et al.*, 1997; Wandji *et al.*, 1998) with demonstrated critical roles in granulosa cell proliferation and inhibition of apoptosis, two processes required for successful follicular maturation (Baker *et al.*, 1996; Baumgarten *et al.*, 2017; Mazerbourg *et al.*, 2003; Spicer and Aad, 2007; Zhou *et al.*, 1997; Peruzzi *et al.*, 1999). *Igf1* and *Igf1r* mRNAs are expressed in the granulosa cells of mouse ovarian follicles (Adashi *et al.*, 1997; Wandji *et al.*, 1998), thus the observed reduction in these mRNAs in this study is likely due to disruptions in this compartment. We hypothesized that reduced ligand and receptor would manifest as reduced IGF1 signaling pathway activity in the ovaries of DBP-treated mice. Although *Igf1r* mRNA expression has been described as constitutive in nearly all stages of development (Wandji *et al.*, 1998), we detected pIGF1R, an indicator of active IGF1R signaling, in all follicle types except primordial with the highest positivity observed in secondary follicles. This pattern of stage-specific positivity was consistent regardless of treatment, in agreement with reports of *Igf1* mRNA expression increasing as folliculogenesis progresses (Wandji *et al.*, 1998). Interestingly, we also observed active IGF1 signaling positivity in atretic follicles, a common observation with another study which localized *Igf1r* mRNA staining in ovaries from another strain of mice (Adashi *et al.*, 1997). Most importantly, we observed that DBP exposure significantly reduced the proportion of primary ovarian follicles with active IGF1 signaling in our mice, a finding that is consistent with our mRNA observations.

Follicular expression of *Igf1* has been shown to increase as folliculogenesis progresses and to decrease as follicular atresia advances in mice (Wandji *et al.*, 1998). Furthermore, in vivo deletion or pharmacological inhibition of IGF1R in mouse granulosa cells leads to increased apoptosis at all follicular stages, and to failed FSH responses in secondary follicles (Baumgarten *et al.*, 2017) and human cumulus cells (Stocco *et al.*, 2017). Given that DBP-treated mice had reduced active IGF1 signaling in their follicles, we hypothesized that these mice would experience alterations in folliculogenesis when compared to vehicle-treated controls. Using the gold standard endpoint for folliculogenesis in rodents, follicle counts, we detected an overall significant decrease in total follicles in mice treated with DBP at 100μg/kg/day. Further differential follicle count analysis led to the discovery that these mice had fewer primordial and primary follicles compared to controls. Interestingly, mice exposed to DBP at 1000 mg/kg/day also had fewer primordial follicles relative to controls. No other follicle counts were different between treatments. Although the specific follicle types involved were unexpected based on our previous work (Sen *et al.*, 2015), the reductions on small follicles observed demonstrate that DBP-induced reductions in IGF1 pathway activity result in a disrupted folliculogenesis phenotype. A well accepted relationship between IGF1 and FSH with phosphoinositide 3-kinase (PI3K) signaling, a significant player in early folliculogenesis (recently reviewed by (Chen *et al.*, 2020)), exists in the ovary. Specifically, FSH-induced activation of AKT, a key kinase downstream of PI3K, in human, mouse, and rat granulosa cells has been shown to require IGF1R activity (Zhou *et al.*, 2013; Baumgarten *et al.*, 2014). Therefore, it is reasonable to suspect that reduced IGF1R activity in the ovaries of our mice disrupts PI3K and AKT activation in small follicles, thus influencing their survival and maturation. The involvement of PI3K in the mechanism of action of other phthalates, DEHP and MEHP, has been proposed to involve excessive recruitment of primordial follicles into the primary stage (Hannon *et al.*, 2014, 2015). While the involvement of PI3K is common between the two studies, our data showing loss of both follicle types strongly suggests a different functional outcome following the interaction of this pathway with each phthalate.

At first glance, the present results seem to contradict our previous report that a 10-day exposure to the same dosages of DBP reduce the number of antral follicles without affecting the smaller follicles (Sen *et al.*, 2015). Interestingly, when the difference in length of exposure is considered (10 days in Sen *et al.* vs 20-32 days in present study), this contrast suggests that continued antral follicle loss over time leads to subsequent loss of follicles at the earlier stages of development. This idea is supported by the lack of significant differences in atretic follicle counts between treatments in this study. Specifically, replacement of lost antral follicles by maturation of smaller follicles may impede the timely observation of high levels of atresia. Alternatively, if the losses are occurring in the small follicle populations only, then atresia would be difficult to identify using standard criteria as these follicles are thought to perish via non-apoptotic pathways (Tingen *et al.*, 2009). Additionally, the ovaries evaluated in the previous study were collected in diestrus while the present study focused on proestrus ovaries. When taking a snapshot of folliculogenesis such as the ones taken in these studies, it is key to recognize that the small and large follicle populations present in the ovary are dynamic throughout the estrous cycle (Mandl and Zuckerman, 1952). Specifically, classical studies indicate that smaller follicles are more abundant in diestrus and that more atretic follicles are detected in metestrus (Butcher and Kirkpatrick-Keller, 1984). Therefore, it is possible that estrous stage specific ovarian sensitivities to chemical insults may exist when follicle counts are examined cross-sectionally. This possibility highlights the need to strategically design folliculogenesis toxicity studies to yield detailed evaluations of ovarian follicle dynamics in the context of DBP and other phthalate exposures. Additionally, understanding the influence of age and the estrous cycle on the ovarian response to DBP exposure are logical next steps in this research.

Based on previous work showing their importance to IGF1R signaling, other molecular events that may be investigated further include DBP influence on the expression and activity of protein tyrosine phosphatases which dephosphorylate IGF1R (Kenner *et al.*, 1996; Maile and Clemmons, 2002b, 2002a) or GATA4 and GATA6 whose activity is implicated in the biological activity of IGF1 in granulosa cells (Bennett *et al.*, 2013). Finally, it has been proposed that the exposure of a follicle to increasing levels of FSH allows it to benefit from locally produced IGF1 and vice-versa, thus, suggesting that the combined actions of these hormones play an essential role in the establishment of dominance (Zhou *et al.*, 2013). In fact, human dominant follicles have been shown to contain higher concentrations of IGF1 and estradiol than non-dominant follicles (Eden *et al.*, 1988). Although conducted using high doses that do not represent human exposures in vivo, a study on rat granulosa cells and follicles revealed that DBP exposure impairs FSH signaling via downregulation of its receptor (Wang *et al.*, 2016). These reports, together with our previous study showing reduced antral follicle numbers in diestrus (Sen *et al.*, 2015), strongly suggest the need to investigate the effects of DBP on follicular gonadotropin sensitivity and selection for ovulation.

Our finding that DBP tampers with the availability of the intraovarian IGF1 system in mice is noteworthy and, although more research is still needed, provides a significant new insight into the molecular mechanisms that respond to phthalate exposure in the ovary. Since insufficient activation of the IGF system has been suggested as a potential explanation for diminished clinical FSH-response (Baumgarten *et al.*, 2014) and reduced reproductive capacity (Baumgarten *et al.*, 2015) in women, our results showing an impaired ovarian IGF system in DBP-treated mice are highly significant. Specifically, they move the field toward a better mechanistic understanding of the associations between high urinary phthalate burden and reduced ovarian reserve (Messerlian *et al.*, 2016) and oocyte retrieval (Hauser *et al.*, 2016) in women undergoing assisted reproduction. Therefore, pursuing epidemiological studies evaluating mediation via IGF system component levels in follicular fluid would be ideal next steps for this research.

## AUTHOR CONTRIBUTIONS

EJ performed experiments, prepared the first draft of the manuscript, and co-wrote subsequent revisions. MM, XL, KM, and KB contributed technically to the experiments and performed manuscript revisions.

ZRC designed the project, performed the animal work, supervised all experimentation and data analysis, and performed final revisions to the manuscript.

## ACKNOWLEDGEMENTS

The authors thank members of the Craig laboratory for their technical assistance.

